# Mechanisms of convergent egg-provisioning in poison frogs

**DOI:** 10.1101/653501

**Authors:** Eva K. Fischer, Alexandre B. Roland, Nora A. Moskowitz, Charles Vidoudez, Ndimbintsoa Ranaivorazo, Elicio E. Tapia, Sunia A. Trauger, Miguel Vences, Luis A. Coloma, Lauren A. O’Connell

**Affiliations:** Department of Biology, Stanford University, Stanford, California, United States of America; Center for Systems Biology, Harvard University, Cambridge, Massachusetts, United States of America; FAS Small Molecule Mass Spectrometry Facility, Harvard University, Cambridge, Massachusetts, United States of America; Department of Biology, Faculty of Science, Antananarivo University, Antananarivo, Madagascar; Centro Jambatu de Investigación y Conservación de Anfibios, Fundación Otonga, Quito, Ecuador; Braunschweig University of Technology, Zoological Institute, Braunschweig, Germany

**Keywords:** parental care, poison frogs, egg provisioning, oxytocin, alkaloids

## Abstract

Parental provisioning of offspring with physiological products occurs in many animals. Within amphibians, maternal provisioning has evolved multiple times, including in South American dendrobatid and Malagasy mantellid poison frogs. In some of these species, mothers feed unfertilized eggs to their developing tadpoles for several months until tadpoles complete metamorphosis. We conducted field studies in Ecuador and Madagascar to ask whether convergence at the behavioral level provides similar benefits to offspring and whether nursing behavior relies on shared neural mechanisms across frogs and vertebrates more broadly. At an ecological level, we found that nursing allows poison frog mothers to provide chemical defenses to their tadpoles in both species. At the level of brain regions, nursing behavior was associated with increased neural activity in the lateral septum and preoptic area in both species, demonstrating recruitment of shared brain regions in the convergent evolution of maternal care within frogs and across vertebrates. In contrast at a molecular level, only mantellids showed increased oxytocin neuron activity akin to that in nursing mammals. Taken together, our findings demonstrate that convergently evolved maternal provisioning behavior provides similar benefits to offspring and relies on similar brain regions. However, the molecular mechanisms underlying the convergence in nursing behavior may be different, suggesting evolutionary versatility in the mechanisms promoting maternal behavior.

## Introduction

Parental care is an important evolutionary innovation, allowing exploitation of novel habitats, influencing fitness and survival of parents and offspring, and serving as an evolutionary precursor to the emergence of other complex social behaviors [1]. Nursing behavior is a specialized form of parental care that is particularly costly as it entails provisioning offspring with resources produced by the parents’ own physiology. Nursing behavior includes lactation in mammals [2], crop milk feeding in birds [3], egg provisioning in amphibians [4], and skin-feeding in fish [5] and caecilians [6]. This remarkable behavior requires coordinated physiological and neural changes, many of which remain poorly understood.

Despite parental behavior being widespread in the animal kingdom, little is known about the neuroendocrine basis of maternal care in non-mammalian species. Accumulating evidence suggests that maternal behavior is governed by shared neural and molecular mechanisms across mammals [7]; however, since maternal behavior coupled with lactation has a single evolutionary origin in mammals, a perspective from outside the mammalian lineage is needed to determine if alternative mechanistic ‘solutions’ can facilitate analogous provisioning behavior, or if the convergent evolution of nursing behavior has occurred via repeated recruitment of similar underlying mechanisms.

Amphibians have independently evolved provisioning behavior many times [8–10], including in some South American (e.g. [4]) and one Malagasy poison frog [11,12]. These species belong to the unrelated families Dendrobatidae and Mantellidae that diverged roughly 140 million years ago and have independently evolved alkaloid-based chemical defenses, warning coloration, and nursing behavior (Figure 1). Nursing behavior in these frogs involves mothers repeatedly visiting offspring to deposit unfertilized trophic (i.e. nutritive) eggs for months until tadpoles complete metamorphosis. The origin of maternal provisioning behavior in poison frogs is tightly linked to ecology: while most poison frogs release tadpoles into large water bodies, nursing species place tadpoles into small volume pools held by terrestrial plants. Thus, the evolution of nursing behavior is associated with colonization of novel habitats where predation and competition are low, but food is scarce [9]. Studies in dendrobatids have established that extended provisioning is costly to mothers, but benefits offspring [9,13], and that tadpoles fed by their mothers carry alkaloids [14,15]. As poison frogs acquire alkaloids from a diet of leaf-litter arthropods to which tadpoles do not have access, this suggests the transfer of chemical defenses from mothers to offspring may be an additional advantage of egg-provisioning.

**Figure 1.**
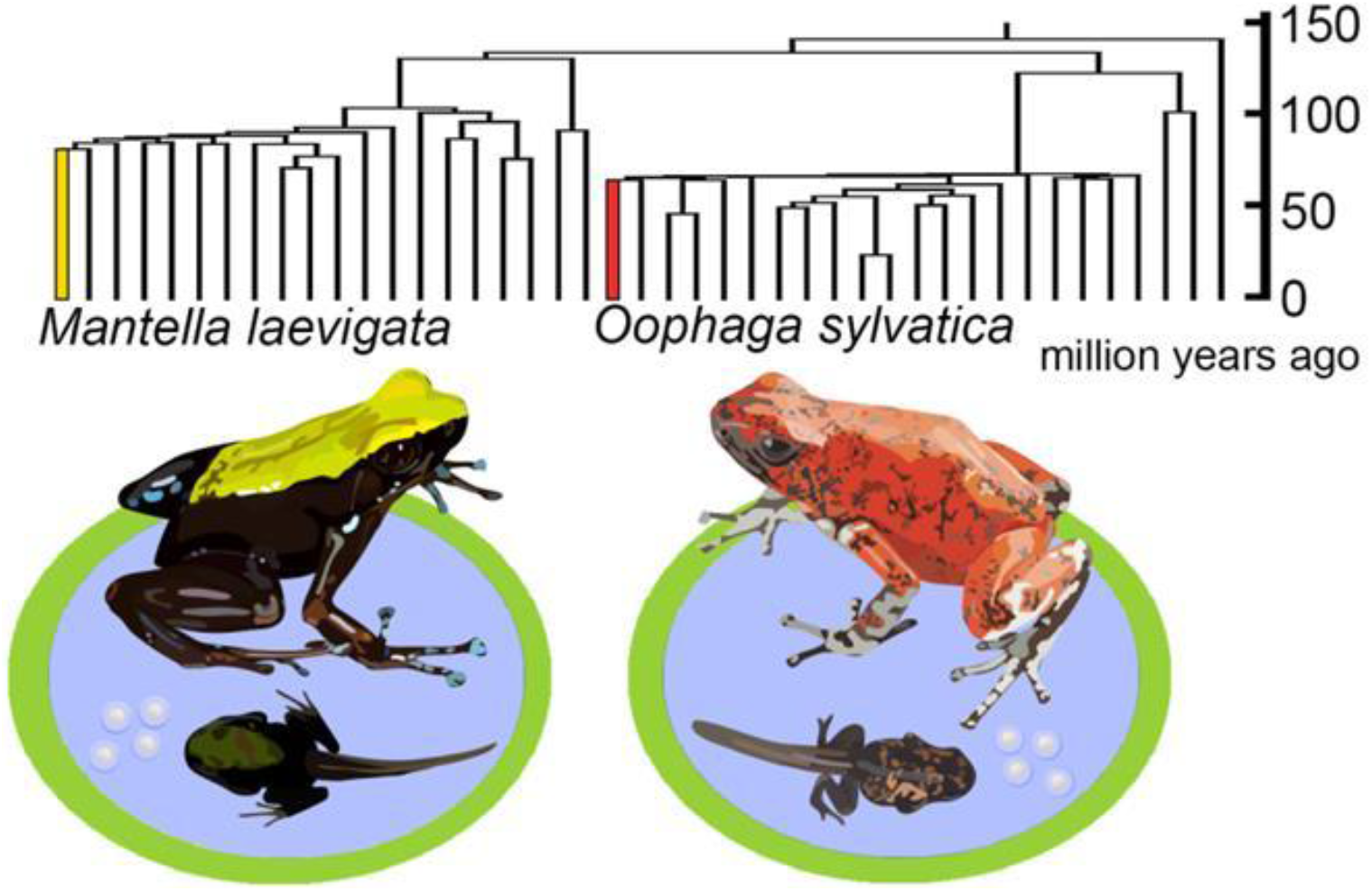
Nursing behavior has evolved independently in amphibians. Malagasy mantellids and South American dendrobatids belong to different major frog clades that diverged roughly 140 million years ago. *Mantella laevigata* (left) and *Oophaga sylvatica* (right) and show striking convergence in a number of traits, including alkaloid sequestration for chemical defense, aposematic coloration, and maternal behavior in the form of egg provisioning to developing tadpoles. Unlabeled branches of the tree are representatives of other anuran families; labeled tree in Supplementary Materials.

We tested the hypothesis that the convergent evolution of nursing behavior across South American and Malagasy poison frog lineages facilitates transfer of chemical defenses to offspring and relies on shared neural mechanisms. We collected field samples from the Little Devil poison frog (*Oophaga sylvatica*) in Ecuador and the Climbing Mantella (*Mantella laevigata*) in Madagascar (Figure 1). We first examined the transfer of alkaloids from mothers to tadpoles across these two species that have convergently evolved acquired chemical defenses and maternal care. We then compared neural activity between nursing and non-nursing females in both species across several candidate brain regions involved in parental and social behavior across vertebrates [16]. Finally, we examined activity specifically in oxytocin neurons in the preoptic area, which are known to mediate maternal behavior and nursing in mammals [17]. Taken together, our findings explore proximate neural mechanisms and ultimate ecological outcomes of convergently evolved maternal egg-provisioning in South American and Malagasy poison frogs.

## Methods

### Field sample collection

*Oophaga sylvatica* were collected at field sites near La Florida, Santo Domingo de los Tsáchilas Province, Ecuador in April and May of 2016. These field sites include naturalistic enclosures populated with *O. sylvatica* frogs that deposit tadpoles in artificial pools. We collected control females from enclosures containing only females to ensure these frogs were non-maternal at the time of capture. *Mantella laevigata* were collected on the island reserve of Nosy Mangabe, Madagascar in November 2015 and July 2016. In Madagascar, frogs were free-roaming making it difficult to control for maternal state. To minimize the likelihood that control females were maternal, we collected controls in the dry season when there were limited pools for reproduction. Furthermore, control females were captured when they were not in the vicinity of a pool and only in areas in which pools were devoid of tadpoles at the time of capture.

The sampling protocol for both species was identical. After identifying a pool containing a tadpole, we waited until females arrived and entered the pool. We left females undisturbed until they exited the pool, upon which we captured them, and verified that they had provisioned the tadpole with eggs. We collected tissue from maternal females 30 min after they entered the pool containing a tadpole and only from those females that laid trophic eggs (N=9 *O. sylvatica*; N=5 *M. laevigata*). Control females were collected at matched times of day (N=8 *O. sylvatica*; N=11 *M. laevigata*). Frogs were anesthetized with application of 20% benzocaine gel to the belly and euthanized by rapid decapitation. Brains were removed and placed in 4% paraformaldehyde for 24 hours followed by storage in 1X phosphate buffered saline (1X PBS). The dorsal skin and oocytes were stored in 1 mL of methanol in glass vials for later alkaloid analysis. Other frog tissues were either preserved in 100% ethanol or RNAlater (Thermo Scientific, Waltham, MA, USA). In addition to adult tissues, we collected skin from tadpoles and trophic eggs that were stored in 1 mL methanol in glass vials. We also collected water from tadpole pools and adjacent tadpole-free pools. Muscle and skeletal tissue were deposited in the amphibian collections of Centro Jambatu de Investigación y Conservación de Anfibios in Quito, Ecuador (*O. sylvatica*) and the Zoology and Animal Biodiversity Department of the University of Antananarivo, Madagascar (*M. laevigata*).

All samples were collected and exported in accordance with Ecuadorian (Collection permits: 005-15-IC-FAU-DNB/MA and 007-2016-IC-FAU-DNB/MA; CITES export permit 16EC000007/VS issued by the Ministerio de Ambiente de Ecuador) and Malagasy (Collection permits: 242/15/MEEMF/SG/DGF/DSAP/SCB and 140/16/MEEMF/SG/DGF/DSAP/SCB; CITES export permit: 1051C-EA12/MG15 and 679C-EA08/MG16 issued by the Direction Générale des Forêts et Direction des Aires Protégées Terrestres [Forestry Branch and Terrestrial Protected Areas Directorate of Madagascar]) laws. The Institutional Animal Care and Use Committee of Harvard University approved all procedures (Protocol 15-03-239).

### Detection of alkaloids by mass spectrometry

Alkaloids were extracted as described in detail elsewhere [43] and briefly summarized below. Prior to extraction, skin samples were weighed with an analytical scale. Trophic eggs and oocytes were processed in a similar manner as skins, except that starting material was not weighed. The entire contents of each sample vial (all tissue and the methanol in which it was stored) were emptied into a sterilized Dounce homogenizer. To ensure the transfer of all materials, the empty vial was rinsed with 1 ml of methanol, which was also added to the homogenizer. We added 25 μg of D3-nicotine (Sigma-Aldrich, St Louis, MO, USA) in methanol to each sample to serve as a standard. Samples were ground with the piston ten times in the homogenizer before being transferred to a glass vial. The homogenizer was rinsed with an additional 1 ml of methanol in order to collect all residual alkaloids, and this methanol was also added to the final glass vial. Samples were stored at −20°C until further processing. Alkaloids from water samples were extracted using Oasis HLB VAC RC 30 mg extraction cartridges (Waters Corporation, Milford, Massachusetts, USA) on a vacuum manifold according to manufacturer instructions, including washes of 5% methanol and elution with 100% methanol. To avoid clogging the cartridges, debris were removed from water samples with a coarse sieve prior to processing.

Alkaloids were analyzed using liquid chromatography / tandem mass spectrometry (LC-MS/MS). Samples were run on a Thermo Q-Exactive Plus and a Phenomenex Gemini C18 3 μm 2.1 × 100 mm column (Torrance, CA, USA). Mobile phase A was composed of water with 0.1% formic acid, and mobile phase B was composed of acetonitrile with 0.1 % formic acid. The flow rate was 0.2 ml/ min. The gradient began with 0 % B for one min, then increased linearly to 100% B at 15 min, and held until 18 min. The column was then re-equilibrated to initial conditions for 3 min before the next sample. Blanks were run at regular intervals to ensure no carry over.

Alkaloids were tentatively identified by comparing this LC-MS/MS data set to a data set obtained by gas chromatography / mass spectrometry (GC/MS) from the same samples and used in a previous study exploring environmental variation and chemical defenses in the same frogs [44]. The alkaloids detected by GC/MS in the previous study were identified using mass spectral data provided in Daly et al. [45]. For the LC-MS/MS data in the present study, a Tracefinder (Tracefinder 4.0, ThermoFisher Scientific) library was created with the accurate mass of all frog alkaloids from the Daly database and used to identify and integrate all potential poison frog alkaloids in the samples. This allowed more sensitive detection of alkaloids in all samples compared to GC/MS and permitted us to trace which potential alkaloids were present in the various sample types. Correlation with the previous dataset for the same frogs allows us to select the mostly likely poison frog alkaloid when several candidates with the same mass were present. Files from LC-MS/MS are available on DataDryad (submission pending).

### Brain immunohistochemistry

Brains were removed from 1X PBS and transferred to 30% sucrose solution for cryoprotection at 4°C for 48 hours, embedded in mounting media (Tissue-Tek® O.C.T. Compound, Electron Microscopy Sciences, Hatfield, PA, USA), and rapidly frozen on dry ice. Brain samples were stored at −80°C until cryosectioning into four coronal series at 14μm. After sectioning, slides were dried for 1–3 hours and stored at −80°C until further processing.

We performed a variety of antibody stains to assess levels of neural activity as well as the overlap between active neurons and cell types of interest. We used an antibody for phosphorylated ribosomes (Phospho-S6 Ribosomal Protein [Ser235/236] Antibody #2211, Cell Signaling, Danvers, MA, USA) as a proxy of neural activation [18]. We followed standard immunohistochemical procedures for 3’,3’-diaminobenzadine (DAB) antibody staining. Briefly, we quenched endogenous peroxidases in 30% sodium hydroxide solution, blocked slides in 5% normal goat serum, and incubated slides in primary antibody (pS6, 1:500 in a 2% normal goat serum solution with 0.3% Triton X-100) overnight at room temperature. The following day, slides were incubated in secondary antibody for 2 hours, followed by avidin-biotin complex (ABC Kit; Vector Labs, Burlingham, CA, USA) solution for 2 hours, and treatment with DAB (Vector Labs) for 2 min. We washed slides with 1X PBS before and between all of the above steps. Finally, slides were rinsed in water, counterstained in cresyl violet, dehydrated in a series of ethanol baths (50%, 75%, 95%, 100%, 100%), and cleared with xylenes prior to cover slipping with Permount Mounting Medium (Fisher Scientific).

We performed fluorescent co-labeling for pS6 and oxytocin on alternate sections of the same brains used for the pS6 staining described above. We followed the same general procedure, but incubated slides in a mix of primary antibodies for pS6 (1:500) and oxytocin (1:5000, MAB5296; Millipore Sigma, Burlington, MA, USA) and a mix of fluorescent secondary antibodies (1:200; Alexafluor 488 (pS6) and 594 (oxytocin)). Following incubation in secondary antibody solution, slides were rinsed in water and immediately cover slipped with Vectashield with DAPI (Vector Labs). Antibody specificity was confirmed by lack of signal after pre-incubating the oxytocin antibody with the mesotocin peptide (Bachem, Torrance, CA, USA), which blocked all signal.

### Microscopy and cell counting

Stained brain sections were photographed at 20x magnification. Brightfield sections were imaged on a Zeiss AxioZoom microscope (Zeiss, Oberkochen, Germany) connected to an ORCA-ER camera (Hamamatsu, San Jose, CA, USA). Fluorescent sections were imaged on a Leica DM4B compound light microscope connected to a fluorescent light source. We quantified labeled cells from photographs using FIJI image analysis software [46]. Cell number was counted in a single hemisphere for each brain region in each section in which that region was visible. For pS6, we measured the area of candidate brain regions and counted all labeled cells within a given region. We quantified cell number in thirteen brain regions that modulate social decision-making across vertebrates [16]: the nucleus accumbens (NAcc), the basolateral nucleus of the stria terminalis (BST), the habenula (H), the lateral septum (LS), the magnocellular preoptic area (Mgv), the medial pallium (Mp; homolog of the mammalian hippocampus), the anterior preoptic area (aPOA), the medial preoptic area (mPOA), the suprachiasmatic nucleus (SC), the striatum (Str), the posterior tuberculum (TP; homolog of the mammalian ventral tegmental area), the ventral hypothalamus (VH), and the ventral pallium (VP). For the pS6 co-stain with oxytocin, we quantified cells only in the preoptic area (both aPOA and mPOA), as this is where oxytocin cell bodies are concentrated in the amphibian brain. We counted oxytocin-positive cells, pS6 positive cells, and the number of pS6-oxytocin co-labeled cells.

### Statistical analysis

We analyzed differences in neural activation associated with egg provisioning behavior using generalized linear mixed models. Behavioral group (nursing vs control), brain region, and their interaction were included as main effects predicting the number of pS6 positive cells using a negative binomial distribution appropriate for count data with unequal variances. Individual was included as a random effect to control for the fact that multiple sections of the same brain region and multiple brain regions were quantified for each frog. In addition, we included brain region area as a covariate to control for body size differences between frogs, size differences between distinct brain regions, and rostral to caudal size variation within single brain regions. We characterized regional differences using Tukey post hoc comparisons adjusted for multiple hypothesis testing.

We tested for differences in the number and activity of oxytocin neurons using generalized linear mixed models. To compare the number of neurons, we included behavioral group as a main effect predicting the number of oxytocin neurons using a negative binomial distribution as before. To analyze activity differences in oxytocin neurons, behavioral group predicted the proportion of pS6 positive oxytocin neurons using a binomial distribution. All statistical analyses were performed separately for each species using SAS Statistical Software v. 9.4; (SAS institute for Advanced Analytics) and plots were made using the base package in R v. 3.5.0 (The R Foundation for Statistical Computing). Raw cell counts are in the Supplemental Materials.

We restricted comparisons of chemical profiles between nursing mothers, tadpoles, internal oocytes, trophic eggs, and tadpole water to those alkaloids present across all provisioning females within each species (Supplemental Materials). For each tadpole, oocyte, and trophic egg sample we calculated the proportion of alkaloids in common with the mother’s skin sample. We then averaged the percent overlap within each species; we are not able to directly compare across species as sample sets were run on different columns. We visualized average percent overlap across all samples using the plotrix v. 3.7-4 in R. We also calculated percent overlap in a pairwise fashion for all tissues. Pairwise overlap was calculated using the proportion of alkaloids shared between an individual tissue group pair to the total number of alkaloids across both tissues. We visualized matrices of pairwise comparisons using the corr.plot v. 0.84 in R.

Finally, we also plotted variation in the level of single toxins. Toxin levels are measured as the integrated area of the major adduct extracted ion chromatogram divided by the area of the internal standard (D3 Nicotine). To make values comparable across species, we normalized these values based on the lowest measurable sample for each species. We plotted log abundance values to facilitate visualization across mothers, tadpoles, oocytes, trophic eggs, and water samples which differ in toxin abundance by orders of magnitude. We note that we retained log of zero values as zero because these are biologically meaningful despite being mathematically undefined. We refer to normalized, log-transformed values simply as ‘relative toxin abundance’ (Supplemental Materials). Boxplots were generated using the base package in R v. 3.5.0 (The R Foundation for Statistical Computing).

## Results

### Convergence in alkaloid provisioning to tadpoles via trophic eggs

We hypothesized that nursing in poison frogs might provide tadpoles with chemical defenses in addition to nutritional benefits [14]. We detected 184 putative alkaloids across all *O. sylvatica* and 317 putative alkaloids across all *M. laevigata*. To facilitate comparisons across individuals with highly variable toxin profiles, we restricted analyses to 32 putative alkaloids shared across all nursing females in *O. sylvatica* and 72 putative alkaloids shared across all nursing *M. laevigata* (Supplemental Materials). Many of these alkaloids were also detected in tadpoles suggesting transfer from mothers to offspring (Figure 2a-b). Maternal alkaloids were also detected in internal oocytes, trophic eggs laid for tadpoles, and in tadpoles’ water pools (Figure 2a-b). Generally, mothers and tadpoles had the highest abundance of alkaloids, followed by the oocytes (internal eggs) and trophic eggs fed to the tadpole (e.g. Figure 2c).

**Figure 2.**
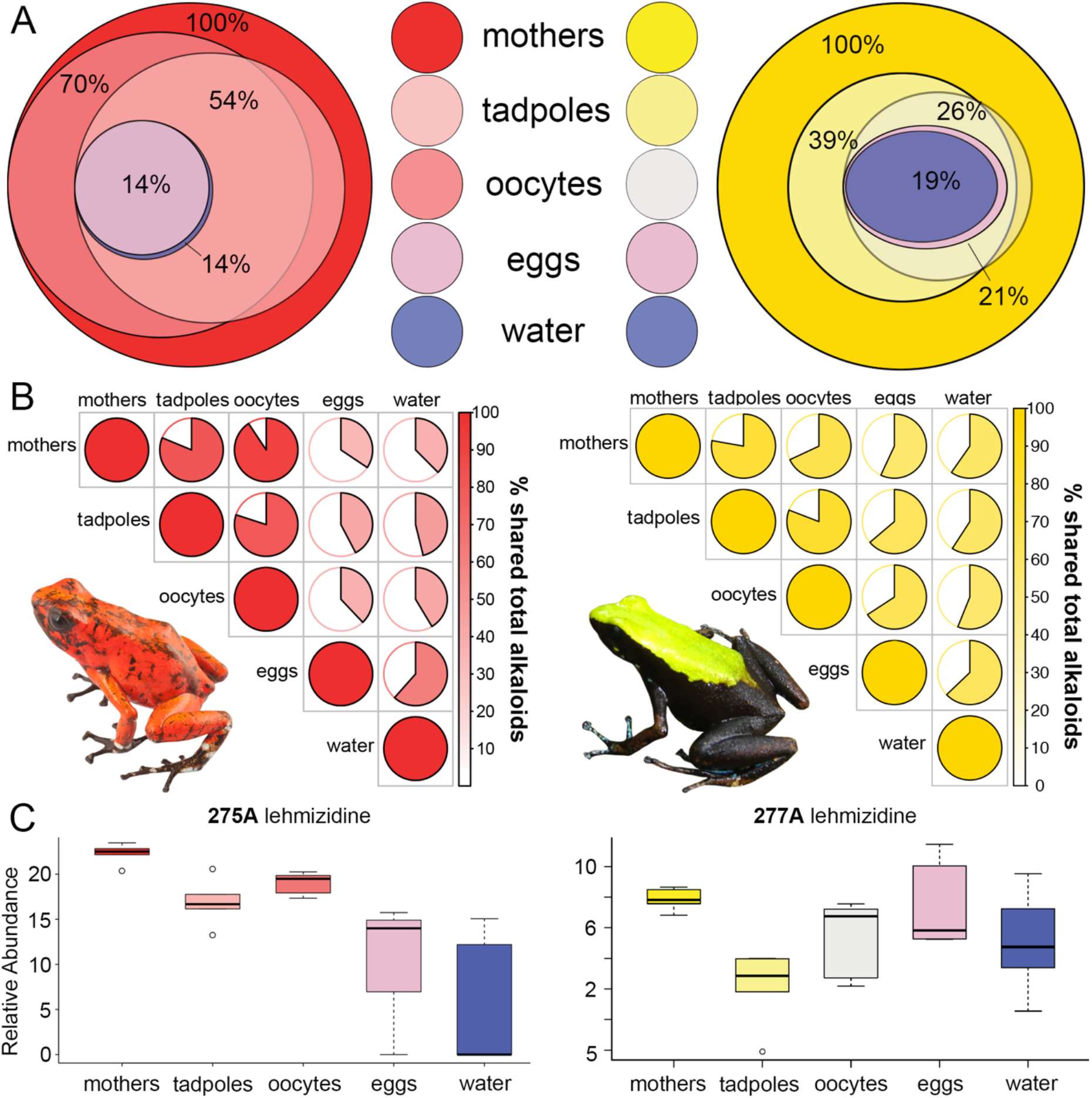
Mothers provision tadpoles with alkaloids across independent lineages of poison frogs. **(A)** Percent overlap of maternal skin alkaloids with alkaloids in tadpole skin, internal oocytes, trophic eggs laid for tadpoles, and water from tadpole deposition sites in *O. sylvatica* (left) and *M. laevigata* (right). The collection of toxins shared across all adult females (32 in *O. sylvatica*, 72 in *M. laevigata*) is considered 100%. Average percentages were calculated from values calculated for each tadpole and tissue paired with its mother. **(B)** Matrices show the percent of shared alkaloids across pairwise sample comparisons for *O. sylvatica* (left) and *M. laevigata* (right) for all toxins across both samples. **(C)** Relative toxin abundances (are measured on an arbitrary scale relative to a 25μg spike of D3 nicotine) for representative lehmizidine alkaloids from *O. sylvatica* (left) and *M. laevigata* (right).

### Shared increases in neural activity in the preoptic area and lateral septum of nursing females across poison frog lineages

To identify brain regions active during egg-provisioning behavior, we compared patterns of neural activity between nursing mothers and non-nursing control females using an immunohistochemical marker for phosphorylated ribosomes (pS6) that serves as a generalized maker of neural activity [18]. We found brain region-specific differences in neural activity in nursing versus non-nursing females in both *O. sylvatica* (group*region: F=7.71, p<0.0001) and *M. laevigata* (group*region: F=3.46, p<0.0001). Two of the brain regions that showed nursing-specific increases in neural activity were shared between species: the medial preoptic area (mPOA; *O. sylvatica*: F=4.56, p=0.033, *M. laevigata*: F=5.63, p=0.018) and the lateral septum (LS; *O. sylvatica*: F=18.77, p<0.0001, *M. laevigata*: F=3.84, p=0.050) (Figure 3). We also found several species-specific activity patterns. Compared to non-nursing controls, nursing *O. sylvatica* had increased activity in the medial pallium (Mp; F=13.52, p=0.0003) and decreased activity in the posterior tuberculum (TP; F=6.78, p=0.009), while nursing *M. laevigata* had increased activity in the nucleus accumbens (NAcc; F=9.10, p=0.003), the basolateral nucleus of the stria terminalis (BST; F=5.36, p=0.021), the anterior preoptic area (aPOA; F=10.19, p=0.0015), the ventromedial hypothalamus (VH; F=14.40, p=0.0002), and the ventral pallium (VP; F=8.80, p=0.0031) (Figure 3). Detailed statistical results are in Supplemental Tables S1 and S2.

**Figure 3.**
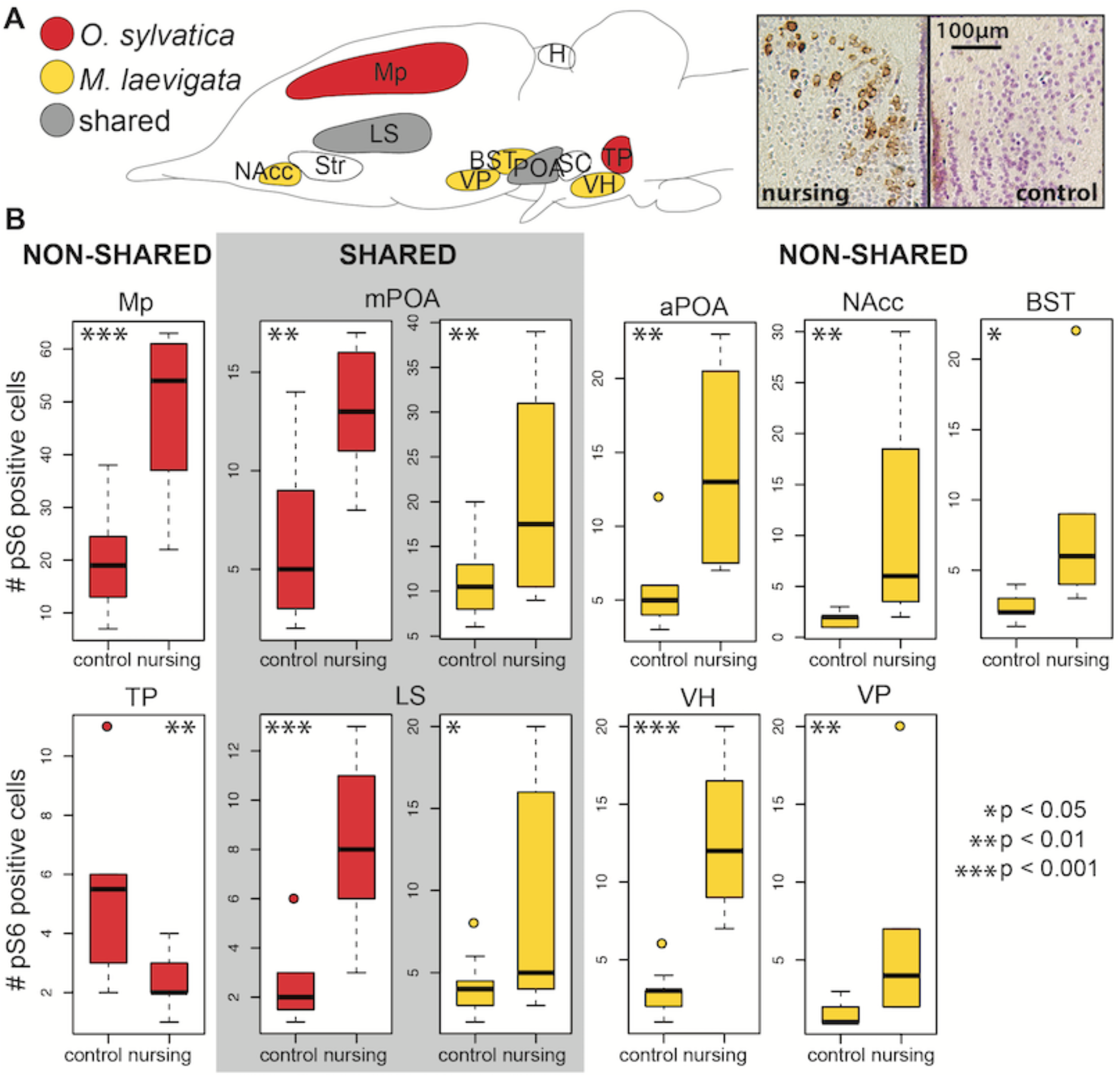
Neural activity differences associated with nursing behavior in independent poison frog lineages. **(A)** Schematic showing brain regions with nursing-associated neural activity patterns in South American *O. sylvatica* (red), Malagasy *M. laevigata* (yellow), and shared across both poison frog clades (grey). Representative micrographs from the mPOA of *M. laevigata* are shown at right. **(B)** Detailed results for shared (center, grey box) and non-shared (left and right) brain regions exhibiting statistically significant activity differences in nursing versus control females. Abbreviations: BST = basolateral nucleus of the stria terminalis, H = habenula, LS = lateral septum, Mp = medial pallium (homolog of the mammalian hippocampus), NAcc = nucleus accumbens, aPOA = anterior preoptic area, mPOA = medial preoptic area, SC = the suprachiasmatic nucleus, Str = striatum, TP = posterior tuberculum, VH = ventral hypothalamus, VP = ventral pallium.

### Opposite patterns of oxytocin neuron activity during nursing behavior in mantellid versus dendrobatid poison frogs

The preoptic area of the hypothalamus is critical for parental behavior across vertebrates [19] and the activation of preoptic area oxytocin neurons appears important for nursing behavior in mammals [17,20]. Behavioral and physiological differences between nursing behavior in mammals and frogs notwithstanding, we examined a link between oxytocin signaling and nursing in poison frogs. To quantify the fraction of oxytocin neurons active during nursing behavior, we fluorescently co-labeled brain tissue for oxytocin and the pS6 marker of neural activity (Figure 4A). In *O. sylvatica*, the proportion of active oxytocin neurons decreased during nursing (F=18.77, p<0.0001), while in *M. laevigata* the proportion of active oxytocin neurons increased during nursing (F=15.55, p=0.0015) (Figure 4B). These opposite patterns were not a byproduct of differences in oxytocin neuron number, as the total number of oxytocin neurons did not differ between behavioral groups in either species (Figure S1).

**Figure 4.**
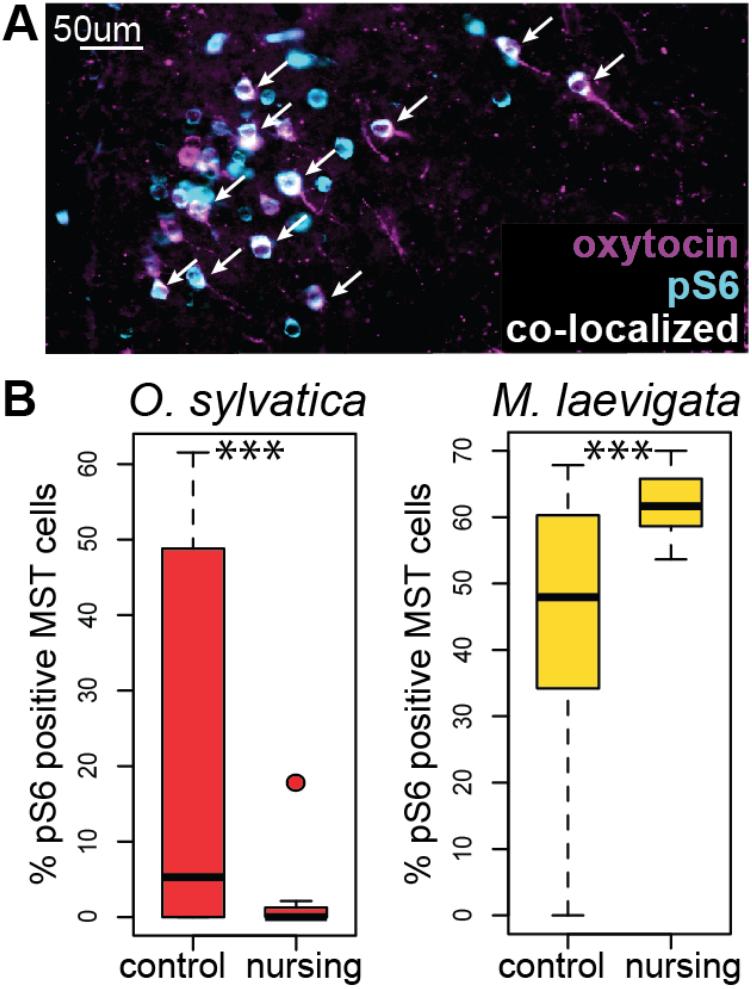
Opposite patterns of oxytocin neuron activity during nursing in dendrobatid and mantellid poison frogs. **(A)** Representative image from the magnocellular preoptic area of *M. laevigata*. Oxytocin neurons are magenta, pS6 positive neurons are cyan, and active oxytocin neurons (i.e. those co-expressing oxytocin and pS6) are white (indicated by arrows). **(B)** The proportion of active oxytocin neurons is decreased during nursing in *O. sylvatica* and increased during nursing in *M. laevigata*.

## Discussion

We explored provisioning of alkaloids to offspring and neuroendocrine mechanisms of nursing behavior across South American and Malagasy poison frogs that have convergently evolved maternal egg-provisioning behavior and chemical defenses. We demonstrated that nursing females in both clades transfer alkaloids to their developing tadpoles and that egg-provisioning behavior relies on partially shared neural mechanisms. We suggest that the transfer of chemical defenses in addition to nutrients provides an additional benefit during the independent emergence and maintenance of this costly maternal behavior in poison frogs belonging to distinct amphibian families.

### Mothers provision tadpoles with chemical defenses

Poison frogs do not synthesize the alkaloids they carry, but rather sequester chemical defenses from their diet of leaf litter arthropods [21], to which developing, aquatic tadpoles do not have access. Although the fertilized egg clutches of several amphibians contain toxins [22], they dissipate during development and tadpoles tend not to carry chemical defenses [23]. However, alkaloid transfer via egg provisioning has been demonstrated in another dendrobatid poison frog, *Oophaga pumilio* [14,15]. Importantly, alkaloids provisioned to these tadpoles offer a survival advantage, where chemically defended tadpoles are less likely to be eaten by predators [24]. We found that both *O. sylvatica* and *M. laevigata* provision their tadpoles with chemical defenses, suggesting convergence in both nutritional and chemical defense provisioning in South American and Malagasy poison frog clades.

While we demonstrated the provisioning of tadpoles with chemical defenses, the predominant mechanism of alkaloid transfer remains uncertain, as alkaloids could be transferred via trophic eggs and/or released from the mother while she is in the water pool and subsequently absorbed by tadpoles. We cannot distinguish between these possibilities at present as we found alkaloids in both tadpole water pools and eggs. However, we have established that oocytes contain alkaloids prior to deposition, highlighting the likely transfer via egg provisioning. The presence of alkaloids in tadpole water pools is an interesting observation that may have implications for tadpole survivorship, as tadpoles only develop granular glands for alkaloid storage during metamorphosis [25]. A toxic water pool may defend pre-metamorphic larvae against predators, competitors, and/or pathogens, though this remains to be tested.

### Shared brain regions associated with maternal provisioning behavior

We found recruitment of shared brain regions in the convergent evolution of nursing behavior in South American and Malagasy poison frogs. In both clades, nursing behavior was associated with increased activity in the lateral septum and preoptic area. The lateral septum modulates goal-directed and social behaviors [26–28]. This broad function encompasses a role in social recognition and memory [29–31], including in the context of paternal care [32]. Although there is little functional information concerning the role of the lateral septum in amphibians, roles for goal-directed motivation and social recognition during nursing are apparent: mothers perform this costly behavior over the course of months, returning at regular intervals to offspring spread across multiple locations. Notably, offspring recognition in poison frogs appears to be based primarily on spatial cues [33,34] providing an opportunity to explore how distinct sensory cues (e.g. spatial cues in frogs versus olfactory and auditory cues in mammals and birds) are integrated by shared neural circuits important in offspring recognition and parental care.

Preoptic area activity is associated with parental care across vertebrates, including in mammals [e.g. 35], birds [36,37], frogs [38], and fish [39]. Recruitment of the preoptic area in the independent evolution of parental care across all major vertebrate lineages suggests that this brain region represents a core node in parental care circuitry independent of sex, species, and care behavior specific phenomena. Indeed, we recently demonstrated sex- and species-independent increases in preoptic area activity during another parental care behavior (tadpole transport) in three species of dendrobatid poison frogs with distinct care strategies (male uniparental, female uniparental, and biparental) [38]. Our findings here expand on this previous observation by demonstrating increased preoptic area activity during a distinct parental behavior (nursing) as well as in a phylogenetically independent clade of frogs (Malagasy mantellids).

### Non-shared brain regions associated with maternal provisioning behavior

In addition to increased neural activity in shared regions, we observed species-specific activity patterns in a number of other brain regions. These differences may be experimental and/or biological in nature, and we emphasize that additional lab-based studies will be necessary to distinguish alternatives. From an experimental perspective, a major difference between our South American and Malagasy field sites was the existence of semi-naturalistic enclosures that allowed us to control social groups in Ecuador but not Madagascar. We collected all non-nursing females when they were alone and not in the immediate proximity of known breeding and tadpole deposition sites. As pS6 activity peaks roughly 45 minutes following behavior, this allows us to exclude the immediate effects of maternal or reproductive behavior, but not possible long-term effects on neural circuit tuning and activity.

At a biological level, we note two key differences in the life history of *O. sylvatica* and *M. laevigata*. First, breeding sites and tadpole deposition sites are distinct in *O. sylvatica*, whereas tadpole deposition sites double as breeding sites in *M. laevigata*. Although we collected females of both species only when they were alone, species level differences in the association between nursing and breeding (i.e. interactions with males) could nonetheless lead to differences in neural activity associated with these behaviors. Second, tadpoles in *O. sylvatica* are obligate egg-feeders that actively beg their mothers for meals by vibrating and nibbling along their legs and abdomen, while *M. laevigata* tadpoles do not appear to exhibit any begging behavior. Thus, species differences in neural activity patterns could be related to differences in interactions among males and females and/or mothers and offspring.

Finally, we note that most of the regions in which we observed species-specific activity patterns belong to the dopaminergic reward system. Homologies of the mesolimbic reward system among vertebrates remain problematic, in particular in amphibians and fish in which data suggest homologs of the mammalian system may be spread across a number of brain regions [40]. Thus, the species-specific neural activity patterns we observe in *O. sylvatica* and *M. laevigata* may represent alternative circuit level ‘solutions’ for motivating goal-directed aspects of maternal care.

### Oxytocin and maternal care

The idea that alternative mechanistic solutions may underlie convergent behavioral phenotypes is further supported by activity patterns of preoptic area oxytocin neurons during nursing. We observed increased oxytocin neuron activity during nursing in *M. laevigata* but decreased oxytocin neuron activity during nursing in *O. sylvatica*, demonstrating contrasting roles for oxytocin signaling during egg-provisioning in poison frogs. Other studies exploring behavioral modulation by oxytocin in frogs give similarly contrasting results, finding increased aggression following oxytocin injections in *Eleutherodactylus coqui* [41], but no effect of oxytocin on parental behavior in the dendrobatid *Ranitomeya imitator* [42]. Taken together, these studies and our findings suggest a nuanced role for oxytocin in amphibian social behavior, with variations linked to species level differences in social behavior and life history. In poison frogs, alternative molecular mechanisms appear to mediate nursing behavior in South American and Malagasy poison frogs, with oxytocin potentially playing a facilitatory role similar to that in mammals in *M. laevigata*, but not *O. sylvatica*.

## Conclusions

Our work demonstrates that convergent nursing behavior in dendrobatid and mantellid poison frog clades facilitates alkaloid provisioning in both clades, and relies on partially shared brain regions but distinct cell types, suggesting that multiple mechanistic solutions may promote vertebrate maternal behavior. While additional work is necessary, we emphasize the value of our comparative design in identifying species level differences and thereby facilitating hypothesis generation and testing with regards to the mechanisms underlying convergent behavioral adaptation. The independent evolution of provisioning behavior across vertebrates provides an exciting opportunity to understand how similar evolutionary advantages of provisioning behaviors with distinct physiological products lead to the targeting of shared versus distinct neural and physiological mechanisms.

## Supporting information

Raw Cell Counts

Toxin Data

## Acknowledgements

We would like to thank María Dolores Guarderas (Wikiri) for support in Ecuador, Joshua Nelson for help with R code, and Deborah Gordon and the members of the O’Connell lab for comments on early versions of this manuscript. We also thank Roberto Marquez for producing the phylogeny in Figure 1. Work in Madagascar was carried out in collaboration with the Malagasy Institut pour la Conservation des Ecosystèmes Tropicaux (MICET) and we are grateful to the Malagasy authorities for issuing research and export permits. Centro Jambatu researchers are indebted to the Ministerio de Ambiente of Ecuador whom support the amphibian research at CJ through the project “Conservation of Ecuadorian amphibian diversity and sustainable use of its genetic resources”.

## Author contributions

LAO conceived of the study, obtained funding, directed the research, and performed the brain immunohistochemistry; LAC and MV contributed to the study design and edited the grants; ABR, EKF, and NR collected samples in Madagascar; EKF, EET, and NAM collected samples in Ecuador; NAM performed the extraction of alkaloids from tissue and water samples; CV and SAT performed the mass spectrometry experiments; ABR performed brain imaging; EKF sectioned brains, performed cell counting, and analyzed cell count data; CV, LAO, NAM, and EKF analyzed the alkaloid mass spectrometry data; EKF and LAO wrote the paper with contributions from all authors.

## References

1. Royle, N.J., Smiseth, P.T., and Kölliker, M. (2012). The Evolution of Parental Care (Oxford University Press).

2. Lefèvre, C.M., Sharp, J.A., and Nicholas, K.R. (2010). Evolution of lactation: ancient origin and extreme adaptations of the lactation system. Annu. Rev. Genomics Hum. Genet. 11, 219–238.

3. Johnston, R.F., and Janiga, M. (1995). Feral Pigeons (Oxford University Press on Demand).

4. Weygoldt, P. (1980). Complex brood care and reproductive behaviour in captive poison-arrow frogs, *Dendrobates pumilio*. Behavioral Ecology and Sociobiology 7, 329–332. Available at: http://dx.doi.org/10.1007/bf00300674.

5. Hildemann, W.H. (1959). A Cichlid Fish, Symphysodon discus, with Unique Nurture Habits. The American Naturalist 93, 27–34. Available at: http://dx.doi.org/10.1086/282054.

6. Kupfer, A., Müller, H., Antoniazzi, M.M., Jared, C., Greven, H., Nussbaum, R.A., and Wilkinson, M. (2006). Parental investment by skin feeding in a caecilian amphibian. Nature 440, 926–929.

7. Dulac, C., O’Connell, L.A., and Wu, Z. (2014). Neural control of maternal and paternal behaviors. Science 345, 765–770.

8. Wells, K.D. (2010). The Ecology and Behavior of Amphibians (University of Chicago Press).

9. Brown, J.L., Morales, V., and Summers, K. (2010). A key ecological trait drove the evolution of biparental care and monogamy in an amphibian. Am. Nat. 175, 436–446.

10. Summers, K. (1999). The effects of cannibalism on Amazonian poison frog egg and tadpole deposition and survivorship in Heliconia axil pools. Oecologia 119, 557–564. Available at: http://dx.doi.org/10.1007/s004420050819.

11. Heying, H.E. (2001). Social and reproductive behaviour in the Madagascan poison frog, *Mantella laevigata*, with comparisons to the dendrobatids. Anim. Behav. 61, 567–577.

12. Brust, D.G. (1993). Maternal Brood Care by *Dendrobates pumilio*: A Frog That Feeds Its Young. J. Herpetol. 27, 96.

13. Dugas, M.B., Moore, M.P., Martin, R.A., Richards-Zawacki, C.L., and Sprehn, C.G. (2016). The pay-offs of maternal care increase as offspring develop, favouring extended provisioning in an egg-feeding frog. J. Evol. Biol. 29, 1977–1985.

14. Stynoski, J.L., Torres-Mendoza, Y., Sasa-Marin, M., and Saporito, R.A. (2014). Evidence of maternal provisioning of alkaloid-based chemical defenses in the strawberry poison frog *Oophaga pumilio*. Ecology 95, 587–593.

15. Saporito, R.A., Russell, M.W., Richards-Zawacki, C.L., and Dugas, M.B. (2019). Experimental evidence for maternal provisioning of alkaloid defenses in a dendrobatid frog. Toxicon. Available at: http://dx.doi.org/10.1016/j.toxicon.2019.02.008.

16. O’Connell, L.A., and Hofmann, H.A. (2011). The Vertebrate mesolimbic reward system and social behavior network: A comparative synthesis. J. Comp. Neurol. 519, 3599–3639.

17. Feldman, R., and Bakermans-Kranenburg, M.J. (2017). Oxytocin: a parenting hormone. Curr. Opin. Psychol. 15, 13–18.

18. Knight, Z.A., Tan, K., Birsoy, K., Schmidt, S., Garrison, J.L., Wysocki, R.W., Emiliano, A., Ekstrand, M.I., and Friedman, J.M. (2012). Molecular profiling of activated neurons by phosphorylated ribosome capture. Cell 151, 1126–1137.

19. Fischer, E.K., Nowicki, J.N., O’Connell, L.A. (2019). Evolution of affiliation: patterns of convergence from genomes to behavior. Phil. Trans. R. Soc. B. 374, 20180242.

20. Russell, J.A., and Leng, G. (1998). Sex, parturition and motherhood without oxytocin? J. Endocrinol. 157, 343–359.

21. Saporito, R.A., Spande, T.F., Martin Garraffo, H., and Donnelly, M.A. (2009). Arthropod Alkaloids in Poison Frogs: A Review of the “Dietary Hypothesis.” Heterocycles 79, 277. Available at: http://dx.doi.org/10.3987/rev-08-sr(d)11.

22. Gunzburger, M.S., and Travis, J. (2005). Critical Literature Review of the Evidence for Unpalatability of Amphibian Eggs and Larvae. J. Herpetol. 39, 547–571.

23. Hayes, R.A., Crossland, M.R., Hagman, M., Capon, R.J., and Shine, R. (2009). Ontogenetic variation in the chemical defenses of cane toads (*Bufo marinus*): toxin profiles and effects on predators. J. Chem. Ecol. 35, 391–399.

24. Stynoski, J.L., Shelton, G., and Stynoski, P. (2014). Maternally derived chemical defences are an effective deterrent against some predators of poison frog tadpoles (*Oophaga pumilio*). Biol. Lett. 10, 20140187–20140187.

25. Stynoski, J.L., and O’Connell, L.A. (2017). Developmental morphology of granular skin glands in pre-metamorphic egg-eating poison frogs. Zoomorphology 136, 219–224.

26. Font, C., Lanuza, E., Martinez-Marcos, A., Hoogland, P.V., and Martinez-Garcia, F. (1998). Septal complex of the telencephalon of lizards: III. Efferent connections and general discussion. J. Comp. Neurol. 401, 525–548.

27. Kondo, Y., Shinoda, A., Yamanouchi, K., and Arai, Y. (1990). Role of septum and preoptic area in regulating masculine and feminine sexual behavior in male rats. Horm. Behav. 24, 421–434.

28. Goodson, J.L., Eibach, R., Sakata, J., and Adkins-Regan, E. (1999). Effect of septal lesions on male song and aggression in the colonial zebra finch (*Taeniopygia guttata*) and the territorial field sparrow (*Spizella pusilla*). Behav. Brain Res. 101, 167–180.

29. Landgraf, R., Gerstberger, R., Montkowski, A., Probst, J.C., Wotjak, C.T., Holsboer, F., and Engelmann, M. (1995). V1 vasopressin receptor antisense oligodeoxynucleotide into septum reduces vasopressin binding, social discrimination abilities, and anxiety-related behavior in rats. J. Neurosci. 15, 4250–4258.

30. Dantzer, R., Koob, G.F., Bluthé, R.M., and Le Moal, M. (1988). Septal vasopressin modulates social memory in male rats. Brain Res. 457, 143–147.

31. Bielsky, I.F., Hu, S.-B., Ren, X., Terwilliger, E.F., and Young, L.J. (2005). The V1a vasopressin receptor is necessary and sufficient for normal social recognition: a gene replacement study. Neuron 47, 503–513.

32. Wang, Z., Ferris, C.F., and De Vries, G.J. (1994). Role of septal vasopressin innervation in paternal behavior in prairie voles (*Microtus ochrogaster*). Proc. Natl. Acad. Sci. U. S. A. 91, 400–404.

33. Stynoski, J.L. (2009). Discrimination of offspring by indirect recognition in an egg-feeding dendrobatid frog, *Oophaga pumilio*. Anim. Behav. 78, 1351–1356.

34. Ringler, E., Pašukonis, A., Ringler, M., and Huber, L. (2016). Sex-specific offspring discrimination reflects respective risks and costs of misdirected care in a poison frog. Anim. Behav. 114, 173–179.

35. Numan, M., and Insel, T.R. (2006). The Neurobiology of Parental Behavior (Springer).

36. Buntin, L., Berghman, L.R., and Buntin, J.D. (2006). Patterns of fos-like immunoreactivity in the brains of parent ring doves (*Streptopelia risoria*) given tactile and nontactile exposure to their young. Behav. Neurosci. 120, 651–664.

37. Ruscio, M.G., and Adkins-Regan, E. (2004). Immediate early gene expression associated with induction of brooding behavior in Japanese quail. Horm. Behav. 46, 19–29.

38. Fischer, E.K., Roland, A.B., Moskowitz, N.A., Tapia, E.E., Summers, K., Coloma, L.A., and O’Connell, L.A. The neural basis of tadpole transport in poison frogs. Available at: http://dx.doi.org/10.1101/630681.

39. O’Connell, L.A., Matthews, B.J., and Hofmann, H.A. (2012). Isotocin regulates paternal care in a monogamous cichlid fish. Horm. Behav. 61, 725–733.

40. Smeets, W.J., Marín, O., and González, A. (2000). Evolution of the basal ganglia: new perspectives through a comparative approach. J. Anat. 196 (Pt 4), 501–517.

41. Ten Eyck, G.R., and Haq, A. ul (2012). Arginine vasotocin activates aggressive calls during paternal care in the Puerto Rican coquí frog, *Eleutherodactylus coqui*. Neurosci. Lett. 525, 152–156.

42. Schulte, L.M., and Summers, K. (2017). Searching for hormonal facilitators: Are vasotocin and mesotocin involved in parental care behaviors in poison frogs? Physiol. Behav. 174, 74–82.

43. McGugan, J.R., Byrd, G.D., Roland, A.B., Caty, S.N., Kabir, N., Tapia, E.E., Trauger, S.A., Coloma, L.A., and O’Connell, L. (2015). Ant and mite diversity drives toxin variation in the Little Devil poison frog. Available at: http://dx.doi.org/10.1101/031849.

44. Moskowitz, N.A., Roland, A.B., Fischer, E.K., Ranaivorazo, N., Vidoudez, C., Aguilar, M.T., Caldera, S.M., Chea, J., Cristus, M.G., Crowdis, J.P., et al. (2018). Seasonal changes in diet and chemical defense in the Climbing Mantella frog (*Mantella laevigata*). PLoS One 13, e0207940.

45. Daly, J.W., Spande, T.F., and Garraffo, H.M. (2005). Alkaloids from amphibian skin: a tabulation of over eight-hundred compounds. J. Nat. Prod. 68, 1556–1575.

46. Schindelin, J., Arganda-Carreras, I., Frise, E., Kaynig, V., Longair, M., Pietzsch, T., Preibisch, S., Rueden, C., Saalfeld, S., Schmid, B., et al. (2012). Fiji: an open-source platform for biological-image analysis. Nat. Methods 9, 676–682.

