## Supplemental Tables & Figures for "Mechanisms of convergent egg-provisioning in poison frogs"

### Supplementary Tables & Figures

**Table S1.** Summary of main statistical effects for neural induction differences.

| | | **df** | **F value** | **p value** |
|---|---|---|---|---|
| ***O. sylvatica*** | group | 1,664 | 1.48 | 0.2240 |
| | region | 12,664 | 13.84 | <0.0001 |
| | group*region | 12,664 | 7.71 | <0.0001 |
| ***M. laevigata*** | group | 1,795 | 6.37 | 0.0118 |
| | region | 12,795 | 17.54 | <0.0001 |
| | group*region | 12,795 | 3.46 | <0.0001 |

**Table S2.** Posthoc statistical results of tests for group differences by region.

| | **region** | **df** | **F value** | **p value** |
|---|---|---|---|---|
| ***O. sylvatica*** | Acc | 1,664 | 0.72 | 0.3958 |
| | BST | 1,664 | 0.02 | 0.8832 |
| | DV/VP | 1,664 | 0.96 | 0.3272 |
| | H | 1,664 | 2.61 | 0.1067 |
| | Ls | 1,664 | 18.77 | **<.0001** |
| | Mgv | 1,664 | 0.40 | 0.5281 |
| | Mp | 1,664 | 13.52 | **0.0003** |
| | aPOA | 1,664 | 0.34 | 0.5580 |
| | mPOA | 1,664 | 4.56 | **0.0331** |
| | SC | 1,664 | 0.09 | 0.7603 |
| | Str | 1,664 | 1.10 | 0.2944 |
| | TP | 1,664 | 6.78 | **0.0094** |
| | VH | 1,664 | 0.00 | 0.9849 |
| ***M. laevigata*** | Acc | 1,795 | 9.10 | **0.0026** |
| | BST | 1,795 | 5.36 | **0.0209** |
| | DV/VP | 1,795 | 8.80 | **0.0031** |
| | H | 1,795 | 0.17 | 0.6815 |
| | Ls | 1,795 | 3.84 | **0.0503** |
| | Mgv | 1,795 | 3.68 | 0.0556 |
| | Mp | 1,795 | 0.00 | 0.9543 |
| | aPOA | 1,795 | 10.19 | **0.0015** |
| | mPOA | 1,795 | 5.63 | **0.0179** |
| | SC | 1,795 | 1.80 | 0.1797 |
| | Str | 1,795 | 3.23 | 0.0728 |
| | TP | 1,795 | 0.88 | 0.3485 |
| | VH | 1,795 | 14.40 | **0.0002** |

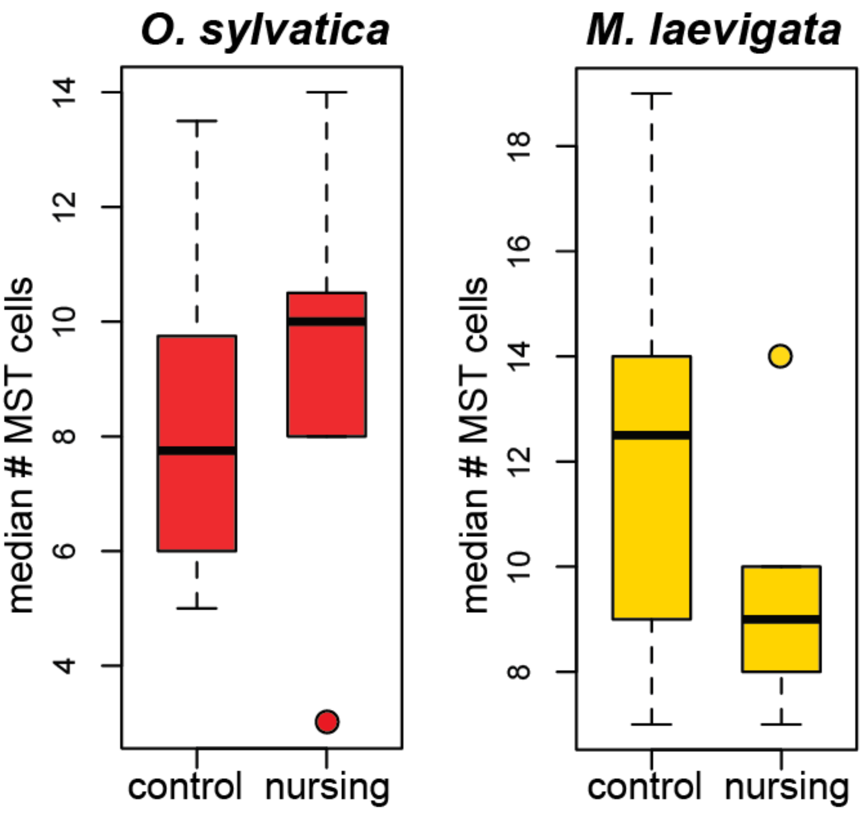


**Supplementary Figure S1. No difference in the number of oxytocin neurons in nursing females. dendrobatid and mantellid poison frogs.** The median number of oxytocin positive cells did not differ between nursing mothers and non-nursing controls in either *O. sylvatica* or *M. laevigata*.

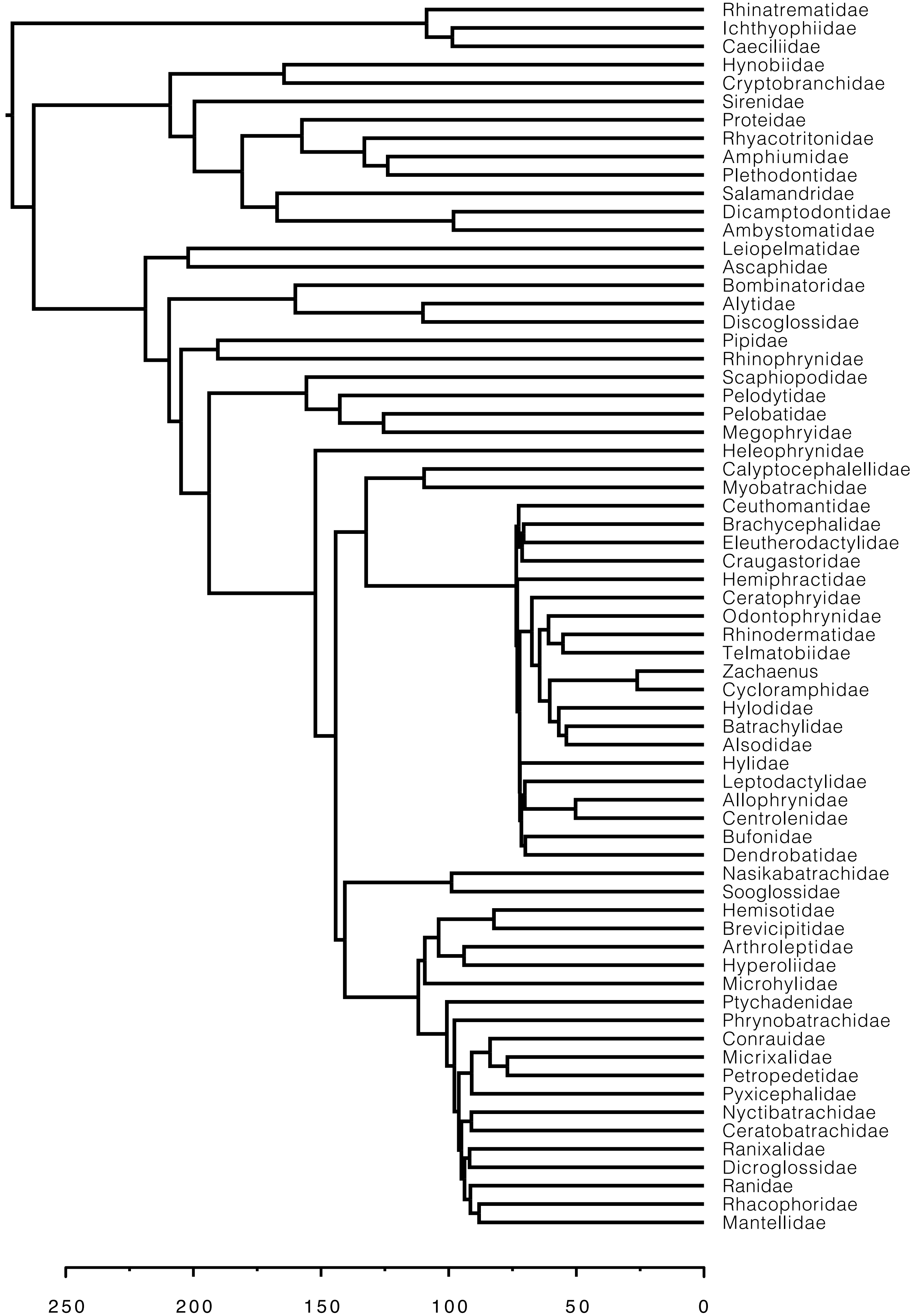

Rhinatrematidae
Ichthyophiidae
Caeciliidae
Hynobiidae
Cryptobranchidae
Sirenidae
Proteidae
Rhyacotritonidae
Amphiumidae
Plethodontidae
Salamandridae
Dicamptodontidae
Ambystomatidae
Leiopelmatidae
Ascaphidae
Bombinatoridae
Alytidae
Discoglossidae
Pipidae
Rhinophrynidae
Scaphiopodidae
Pelodytidae
Pelobatidae
Megophryidae
Heleophrynidae
Calyptocephalellidae
Myobatrachidae
Ceuthomantidae
Brachycephalidae
Eleutherodactylidae
Craugastoridae
Hemiphractidae
Ceratophryidae
Odontophrynidae
Rhinodermatidae
Telmatobiidae
Zachaenus
Cycloramphidae
Hylodidae
Batrachylidae
Alsodidae
Hylidae
Leptodactylidae
Allophrynidae
Centrolenidae
Bufonidae
Dendrobatidae
Nasikabatrachidae
Sooglossidae
Hemisotidae
Brevicipitidae
Arthroleptidae
Hyperoliidae
Microhylidae
Ptychadenidae
Phrynobatrachidae
Conrauidae
Micrixalidae
Petropedetidae
Pyxicephalidae
Nyctibatrachidae
Ceratobatrachidae
Ranixalidae
Dicroglossidae
Ranidae
Rhacophoridae
Mantellidae
250
200
150
100
50
0

**Supplementary Figure S2.** The phylogeny used in Figure 1 was pruned from the anuran tree in Pyron (2014), which originally included 3309 species. One species per 64 families was arbitrarily chosen using the drop.tip() function in the R package ape. Reference: Pyron, R.A. (2014). Biogeographic Analysis Reveals Ancient Continental Vicariance and Recent Oceanic Dispersal in Amphibians. System. Biol. *63*(5), 779-797.
